# SimSearch: A Human-in-the-Loop Learning Framework for Fast Detection of Regions of Interest in Microscopy Images

**DOI:** 10.1101/2022.04.05.487117

**Authors:** Ankit Gupta, Alan Sabirsh, Carolina Wählby, Ida-Maria Sintorn

**Affiliations:** Department of Information Technology, Uppsala University, Uppsala, Sweden; Advanced Drug Delivery, Pharmaceutical Sciences, R&D, Astrazeneca, Mölndal, Sweden; Science for Life Laboratory, 75237, Sweden; Vironova AB, 11330 Stockholm, Sweden

## Abstract

**Objective:** Large-scale microscopy-based experiments often result in images with rich but sparse information content. An experienced microscopist can visually identify regions of interest (ROIs), but this becomes a cumbersome task with large datasets. Here we present SimSearch, a framework for quick and easy user-guided training of a deep neural model aimed at fast detection of ROIs in large-scale microscopy experiments.

**Methods:** The user manually selects a small number of patches representing different classes of ROIs. This is followed by feature extraction using a pre-trained deep-learning model, and interactive patch selection pruning, resulting in a smaller set of *clean* (user approved) and larger set of *noisy* (unapproved) training patches of ROIs and background. The pre-trained deep-learning model is thereafter first trained on the large set of *noisy* patches, followed by refined training using the *clean* patches.

**Results:** The framework is evaluated on fluorescence microscopy images from a large-scale drug screening experiment, brightfield images of immunohistochemistry-stained patient tissue samples, and malaria-infected human blood smears, as well as transmission electron microscopy images of cell sections. Compared to state-of-the-art and manual/visual assessment, the results show similar performance with maximal flexibility and minimal a priori information and user interaction.

**Conclusions:** SimSearch quickly adapts to different data sets, which demonstrates the potential to speed up many microscopy-based experiments based on a small amount of user interaction.

**Significance:** SimSearch can help biologists quickly extract informative regions and perform analyses on large datasets helping increase the throughput in a microscopy experiment.

## 1 Introduction

Microscopy imaging is an essential tool for investigating complex biological processes. With the increase in high throughput imaging techniques, the amount of data being generated is increasing at an unprecedented rate. The high resolution images created in biology often contain rich but sparse information, i.e., only a small section of the data contains information useful for deeper analysis. Expert annotators are usually required for sifting through the data and generating insights about those informative regions. As a result, manual annotation is difficult to scale with the amount of data being generated. Hence, there is a need for an efficient and flexible user-guided detection of informative regions in the data.

### 1.1 Related Work

Current state-of-the-art tools (1; 2; 3; 4; 5) in biomedical image analysis rely on manual feature selection and training machine learning classifiers with predefined extracted features. In CellProfiler (3), the user is asked to create a ‘pipeline’ consisting of a sequence of individual modules typically performing image processing, object segmentation and feature measurements. For each module, methods need to be chosen and parameters interactively tuned. The extracted measurements can then be imported into CellProfiler Analyst (4) to classify different objects based on a few user-defined examples and machine learning classifiers such as SVM and RandomForest.

Ilastik (5) provides a graphical user interface (GUI) for semantic image segmentation where the user defines training regions using mouse-clicks and brush strokes. Pixel classification is based on a set of user selected pre-defined local neighborhood features, such as color, intensity, and texture, on which the classifier makes its predictions.

In both these tools, the user is expected to select the image features required to classify the ROIs which means that the results are limited by the ability of selected features to separate the ROIs of different classes. Any shift in imaging conditions, intensity or texture will easily cause the features to fail and, hence, reduce the performance of these tools (5). Furthermore, even though both tools provide estimators of feature importance to prune irrelevant features, method choices and feature selection still require expertise in image analysis and biology.

Deep learning techniques have proven to be very effective and useful for biological image analysis (6; 7). In many situations, these techniques provide a distinct advantage over task-specific feature engineering which is time-consuming and often inadequate for dealing with contextual variations, and instead learn hierarchical feature representations of the image content from labeled samples (8). Although the learned features are task and data specific, the first few layers of the representations are often generic (edges, blobs, shapes) and can serve as features to represent different ROI classes as shown in this paper.

However, deep learning requires a large amount of labeled data to perform well, which in case of biological data is very resource demanding to produce due to its requirement of domain-expertise that increases the logistic and financial burden. Recently, the emergence of self-supervised learning techniques (9), (10) has shown great promise in reducing the requirement of massive amounts of labeled data. In self-supervised learning, the model learns the feature representation of the input data using a pre-designed task without using the annotations. The model can then be finetuned to be used in a variety of down-stream tasks with a small amount of labeled data.

### 1.2 Contribution

In this paper, we introduce a human-in-the-loop ROI detection framework, SimSearch, which combines established deep learning techniques with modified selfsupervised learning to make efficient use of limited user interaction. The main contributions of this paper are as follows:

- An intuitive and interactive GUI framework for ROI detection requiring the user input only on a small number of images in a microscopy dataset.
- The user annotates the images to create a small set of *clean* ROIs consisting of *positive* and *negative* ROIs.
- Using user-input efficiently to create a larger set of *noisy* ROIs from the rest of the dataset and learning the feature representation of the dataset using self-supervised learning.
- Fine-tuning the self-supervised model on the *clean* data with supervised contrastive loss utilizing the user-selected *negative* ROIs in a novel way.

It first uses *prototype* ROIs marked by the user in a few displayed images to extract features from a standard pre-trained deep learning model. Based on the extracted features, similar patches are suggested on the displayed images and the user chooses a similarity threshold using a slider for true *positive* patches and curates the positive set by interactively removing the falsely suggested patches. These interactively removed patches are the *negative* patches. The chosen threshold is used to extract a larger set of *noisy* patches from the images not displayed to the user. These are used to retrain the initial deep learning model in a self-supervised manner to learn the general behaviour of the imaging modality and ROIs. Finally the model is fine-tuned on the curated clean positive and negative patches, resulting in a deep learning model which specifically caters to the task at hand.

The paper is organized as follows. The components of the framework and model training are described in Section 2. Section 3 describes the datasets and evaluation criteria used to demonstrate and evaluate the framework. Section 4 presents the results and discussion of SimSearch on on experiments and an ablation studies, and 5 concludes our findings.

## 2 Methods

### 2.1 Overview and Notations

The framework pipeline consists of data preprocessing, combined manual and automated iterative patch selection, deep learning model training, and final patch confidence thresholding. All the steps are described in detail in the following sections and illustrated in Figure 1. Since the input may come from different modalities with different bit-depth and number of channels, all available images are pre-processed and the user is asked to select relevant channels to include in the ROI detection. In fact, irrelevant image channels may introduce unwanted bias in ROI selection, and focusing on relevant channels reduces computation time and latency. Next, the user initiates the iterative patch selection process on a displayed subset of the images. The user selects ROIs by drawing rectangular patches in the images. These patches are denoted as the *prototype* patches of the instantiated class. The corresponding feature vectors extracted by a simple pre-trained model are called *prototype vectors*. New patches in the displayed images are suggested to the user based on their similarity (see Section 2.3.1) to the prototype patches. The user chooses a similarity threshold using a slider, manually curates the suggestions and then approves *clean* patches consisting of the *positive* and *negative* (the manually curated) ROIs for each class, i.e., belonging to the class, and not belonging to the class respectively. Based on the similarity threshold, *noisy* patches, i.e., unapproved and uncurated ROIs are generated from the rest (undisplayed) of the images.

**Figure 1:**
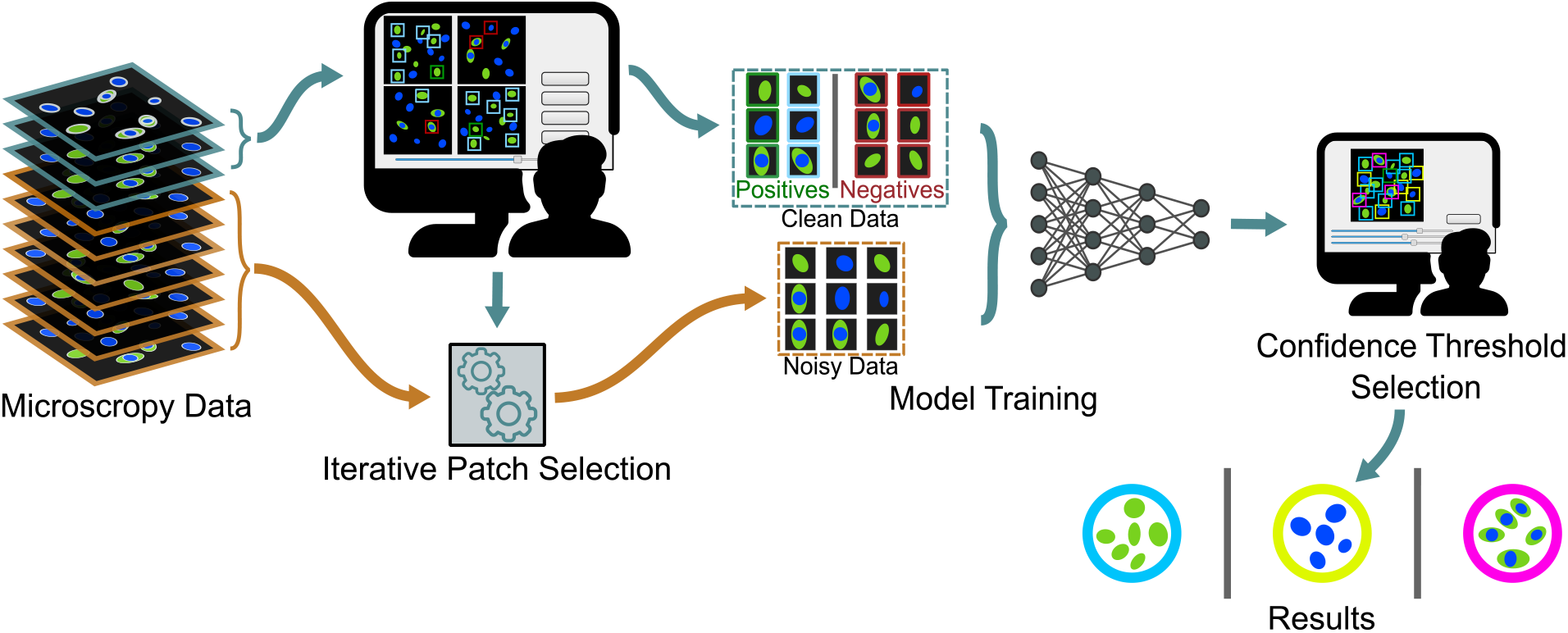
Overview of the SimSearch framework. The human-in-the-loop system requires minimal input at each interactive step in the pipeline. Iterative Patch Selection (IPS) is done on a subset of images and the parameters are used to generate noisy data. Thereafter, a deep learning model is trained, and the output is shown to the user in Confidence Threshold Selection. Finally, the model and results are saved.

A task-specific deep learning model is trained next using the *clean* and *noisy* data. The user is then presented with the patch classification result from the trained model, and asked to choose a suitable confidence threshold for the final ROI definition. Finally, the complete dataset is (again) processed by the model, and the ROIs are displayed and/or saved.

### 2.2 Preprocessing and Channel Selection

Images captured by a digital camera connected to a microscope usually have a bit-depth of 12-16 bits/pixel. This wide dynamic range is sometimes useful for precise feature extraction, but the information contained within a smaller dynamic range is often sufficient for detection of ROIs. In addition, fluorescence microscopy images often consist of multiple channels that contain information about different structures. Channels also may be correlated, and in many cases only a subset of the channels are relevant for ROI selection. Hence, to reduce computational costs and speed up the ROI selection, the dynamic range is re-scaled and channel correlation is measured.

All images of a channel are re-scaled to the same dynamic range. We assume that the instrument settings are kept constant throughout the experiment and apply the same gentle normalization to all images in the set. For each channel, we extract the (5, 95) intensity percentile of all images. We then find the (0.5, 99.5) of these intensity percentiles and re-scale intensities in this dynamic range to [0-1]. This minimizes the influence of image artifacts. The mean of the correlation between image channels is calculated and shown to the user for manual channel selection in the GUI. The user can then make a statistically informed decision by clicking on the channels that are relevant for the experiment. The manual channel selection can be skipped if information from all channels should be used for the ROI detection.

### 2.3 Iterative Patch Selection (IPS)

After preprocessing and channel selection, the user is presented with a graphical user interface (GUI) shown in Figure 1 displaying *n* images randomly chosen or selected from the dataset. The user then selects a small number of prototype patches and interacts with the GUI to generate *clean* training patches representing ROIs and background following the flowchart in Figure 2. The number of images, *n*, can be chosen by the user and should preferably be kept small (less manual work) while still contain representative objects and the variation in the dataset.

**Figure 2:**
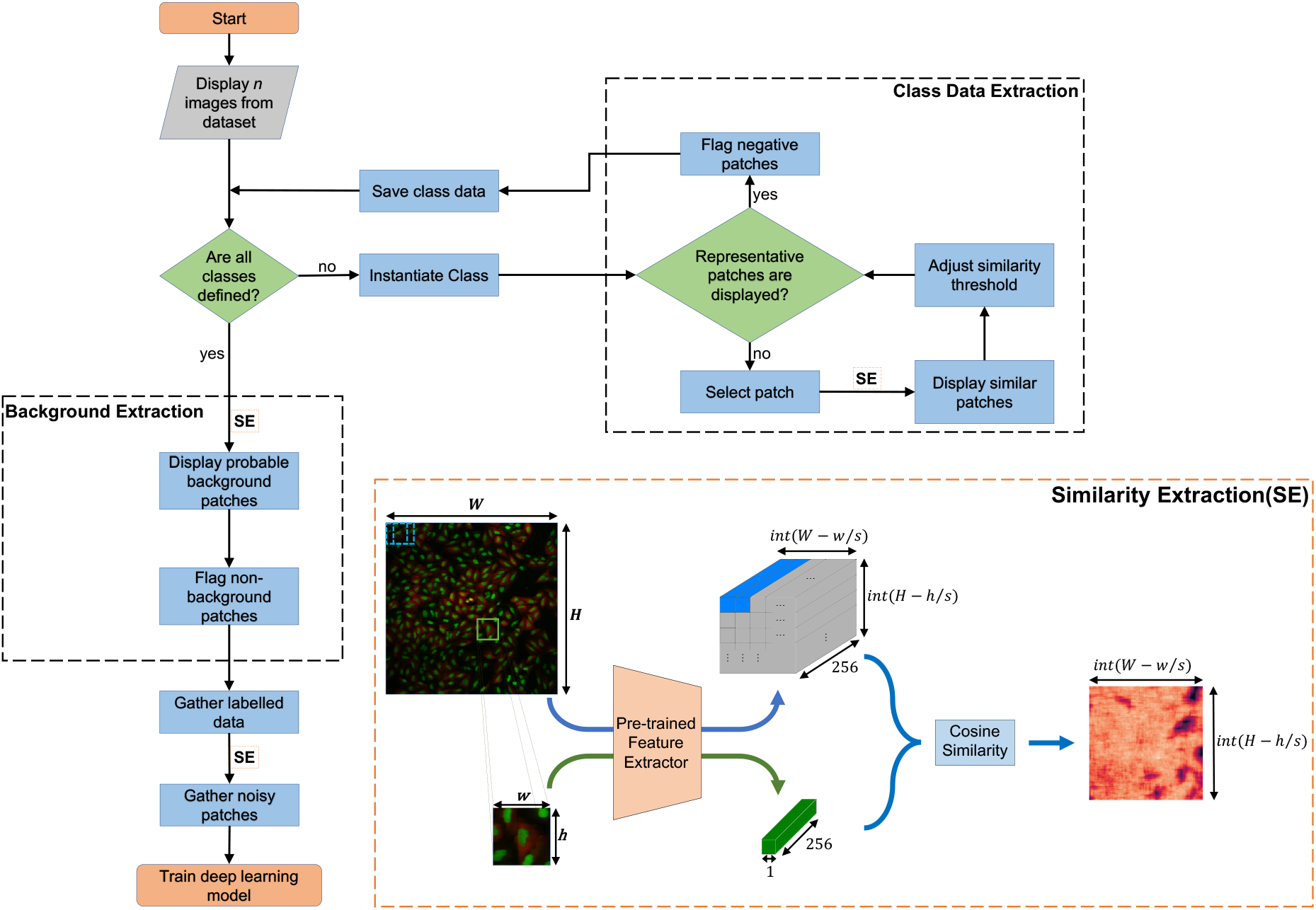
Flowchart of iterative patch selection and the description of Similarity Extraction (SE) (in orange box). The user instantiates classes representing ROIs by selecting patches and creates data following the Class Data Extraction block. The SE is called for feature extraction and detection of similar patches in multiple places in the flowchart. After all the classes are defined, the user marks the background class in the Background Extraction block. Noisy training patches are generated using the class *prototypes* and similarity threshold.

To start the definition of ROIs, the user first instantiates a ROI class and then marks *prototype* patches representing this ROI class. Based on ‘Similarity Extraction’, as described below in Section 2.3.1, patches similar to the prototypes are highlighted in the displayed images. The user then adjusts a similarity threshold to include or exclude more patches based on their similarity to the *prototype* patch. The user can select more *prototype* patches and adjust the threshold to annotate the images until a representative number of patches (typically 80-90% of the objects of the instantiated class in the displayed images) are marked. If incorrect patches are included after thresholding, the user may manually mark them to exclude these patches, which are kept as *negatives*.

Once patches from all the instantiated ROI classes are annotated, the user is presented with patches that are assumed to belong to the background. The user is asked to mark the patches that do not belong to the background. Once this process is completed, all *clean* patches, i.e., positive and negative patches approved by the user, are gathered and the similarity threshold is saved for each ROI class. Thereafter, based on the *prototype* patches and the similarity threshold, *noisy* training patches are extracted from the rest of the dataset.

#### 2.3.1 Similarity Extraction (SE)

With the selection of the first *prototype* patch for a ROI class, the patch shape and stride (0.1 × the patch width) for the experiment are fixed and similar patches are shown to the user following Similarity Extraction as described in Figure 2. The *prototype* and its rotations (90°,180°and 270°) are passed through a pre-trained CNN feature extractor (shown in Figure 3) to get the feature representation of the patch, referred to as *prototype vectors* of that class. In this paper, a deep learning model pretrained on ImageNet(11) dataset is used. This may not be optimal, but sufficient, as the goal of the IPS is to find a few general features roughly representing the ROIs. The primary interest here is to swiftly label many patches.

**Figure 3:**
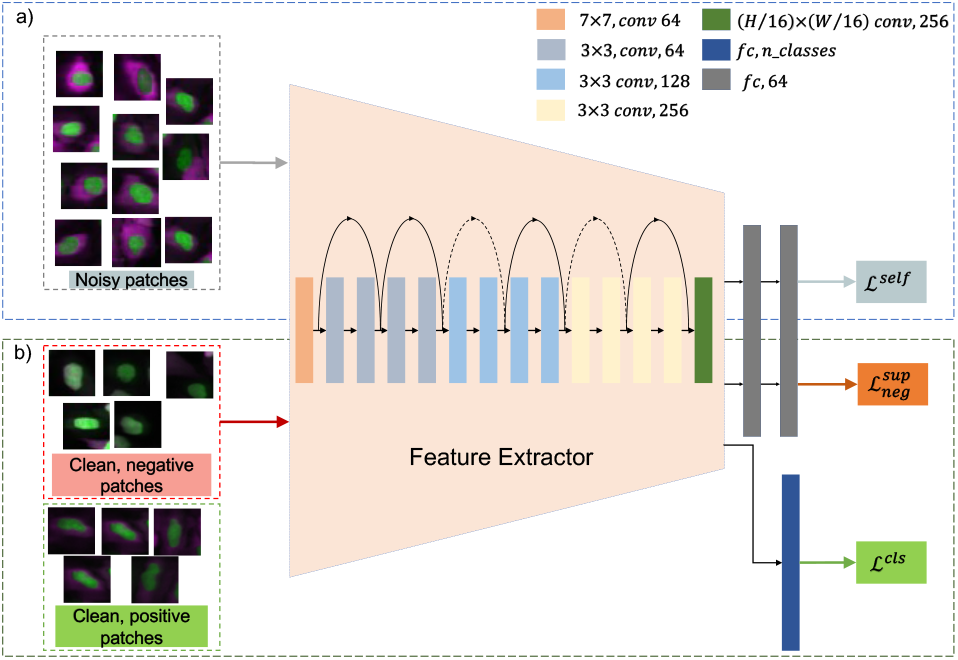
Description of the model architecture and training strategy. Training is divided into two parts. The model is first trained with *noisy* patches generated in 2.3.3 (top). Afterwards, the model is trained with both *clean negative* patches and *clean positive* patches and finally, a fully connected layer is added and trained in the end to make predictions (bottom).

Similarly, the feature representations of the images are extracted by dividing the image into patches using a sliding window with the set patch shape (from the first manually drawn ROI) and stride. The cosine similarity between the *prototypes* and the feature representation of all image patches is calculated, providing a heat-map of similarities over the images. The locations of local maxima above the set similarity threshold from the heat map are calculated and projected back to the image and shown as similar patches to the user. The user can then adjust the threshold to see patches with higher or lower similarity.

However, as the threshold is decreased, all patches displayed might not belong to the instantiated class. This “false-positive” number can be reduced by increasing the threshold but at the cost of getting fewer *clean* positive patches to train the model with. Reducing the threshold would result in more examples but also more false-positive examples. To eliminate this issue, a *negative* flagging step is introduced in the class data selection process. After having selected a similarity threshold, the user is asked to flag the patches that do not belong to the class. This results in more *clean positive* examples to be available for training and also *clean negative* examples to make the classifier more robust towards difficult examples.

All the *prototype* vectors and final similarity threshold are saved once the user is done with *negative* flagging. These are used to extract the background patches, and generate *noisy* samples for the rest of the data, as described in Section 2.3.2 and Section 2.3.3 respectively.

#### 2.3.2 Background Extraction

After labeling all the relevant classes, the user is presented with some previously unlabeled patches that potentially belong to the background and is asked to flag the patches that do not belong to the background. The semi-automated background extraction step is introduced to eliminate the need for manual background patch selection. The patches presented to the user are the unlabeled patches with highest and lowest similarity to the average of all *prototypes* for all classes. The patches with the lowest similarity represent the easy examples of the background and the highest similarity represent difficult examples. An approach similar to the SE is used (see Algorithm 1).

##### Algorithm 1: Find 2*n* probable background patches in the image

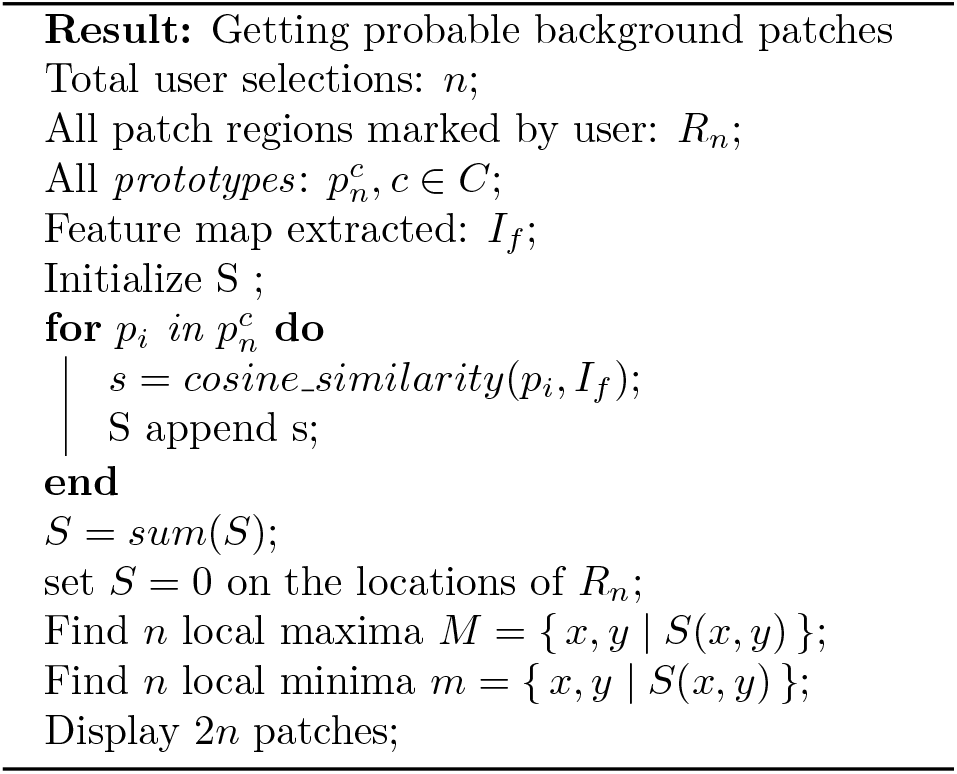

#### 2.3.3 Generation of Noisy Training Data

Based on the prototypes and similarity thresholds (ROIs and background), *noisy* data can be generated automatically from the images in the dataset not displayed to the user. The images are passed through the feature extractor and the local maxima are extracted from the cosine similarity between the feature map and *prototypes*, and thresholded based on the user-selected similarity threshold for the class as described in 2.3.1. The background patches are then similarly generated using the algorithm 1. The generated data is *noisy* due the lack of user verification, but it provides valuable information for tuning the feature extractor to the characteristics of the images and task at hand.

### 2.4 Deep Learning Model Training

The model training is done in two steps, first the noisy patches are used to train the network in selfsupervised way to learn the general representation of the data and, second, the model is refined towards the class representation using the patches verified by the user. Finally, a fully connected layer is added and a classifier is trained.

#### 2.4.1 Training on Noisy Patches

The feature representations of the patches in the datasets are learned on the *noisy* patches using Sim-CLR, the self-supervised contrastive learning framework introduced in (12). Since only the most relevant noisy patches are extracted by the method described in 2.3.3, the feature representation learned by contrastive learning enhances supervised training on the limited amount of labeled data. In SimCLR, a projection head (a multi-layer perceptron (MLP) with one hidden layer) is attached to the pre-trained feature extractor model and the whole architecture is then trained with the *NT-Xent* (normalized temperature-scaled cross entropy) loss using two differently augmented views of the same noisy patch in each minibatch.

The NT-Xent loss for the correlated views of the same example is defined as follows,

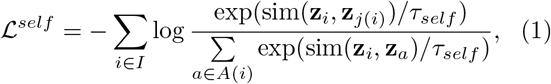

Here, *i* is a sample in a multi-viewed minibatch of size 2*N* where each sample appears twice, augmented differently, i.e., *I* ≡ 1…2*N*. Index *i* is referred to as *anchor*, and *j*(*i*) is called its *positive* and other 2(*N* — 1) indices in *A*(*i*) ≡ *I* \ {*i*} are called *negatives.*Further, sim(**u, v**) =**u**^*T*^**v**/∥**u**∥∥**v**∥ denotes the cosine similarity between *u* and *v* and *τ_self_* denotes the temperature parameter for the self-supervised training.

The initial layers of the feature extractor are frozen to enable only the task specific high-level features to be learned during the noisy refinement. An alternative noisy training scheme has been proposed in (13) which tried to achieve learning with noisy samples by simultaneously training with noisy labels and corrected labels. This scheme was tested but did not lead to improvements since the feature representation of *negative* and *positive* patches could not be separated meaningfully.

#### 2.4.2 Further Refinement on *Clean* Patches

As previously shown in (14), the training procedure above can be generalized to be used in supervised learning conditions where labels of the samples are available. The loss function is then modified to incorporate the class labels for the patches in order for the model to learn the representation that maximizes the similarity between patches of a class (positives) and minimize the similarity to patches of a different class (negatives).

The supervised contrastive loss can be defined as follows,

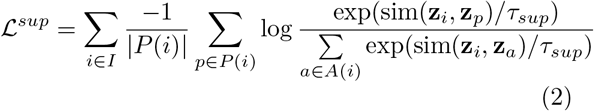

where *A*(*i*) ≡ {*a* | *a* ∈ *I* \ *i*} and *P*(*i*) ≡ {*p* ∈ *A*(*i*): *y_p_* = *y_i_*}.

The *negative* patches marked by the user provide challenging examples where the original similarity measure (from the unrefined model) failed in the patch selection process. We can utilize these *negatives* and add them directly into the denominator of the log term in Eq. 2. The denominator can then be written as,

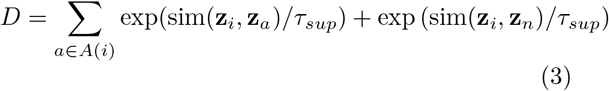

where *A*(*i*) ≡ {{*a*, *n*} | *a* ∈ *I* \ *i,n* ∈ *N*(*i*)} and *N*(*i*) is a set of all *negative* examples marked by the user during the patch selection of class *y_i_*. The final loss then becomes

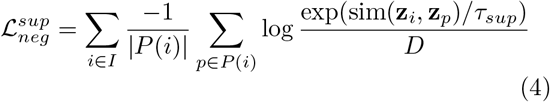

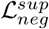 helps the model learn feature representation of the dataset that maximizes the similarity between the *positive* samples of the same class, but minimizes the similarity of *positive* samples of a class with the *negative* samples of that class, and the samples of other classes.

Finally, the projection head is removed and a fully-connected layer is added, see Figure 3. The refined feature extractor part of the model is then frozen and only the fully connected layer is trained with crossentropy loss to get the patch class prediction as output.

Since the data from the user selection tends to be imbalanced, a weighted oversampling was used in both the *noisy* and *clean* patch training steps to prevent overfitting and skewing the results towards the over-represented class. By default, the framework trains the model with both the *noisy* and *clean* patches but noisy training and negative training can be switched off depending on the precision and computation requirements by the user.

### 2.5 Confidence Threshold Selection

The inference is done in a patch-wise manner where the image is divided into patches using a sliding window and a softmax value of the prediction for each class is obtained for the patches. The confidence threshold is then defined as the softmax threshold above which the local maxima in the softmax image are chosen. Results on the dataset images are displayed in the GUI and the user has the option of changing the confidence of each class separately and choosing a threshold that produces the desired result. This gives the user an option to prioritize either precision or recall in accordance with the research question. A higher threshold will result in predictions with higher precision but lower recall and a lower threshold will result in predictions with higher recall and lower precision. After threshold selection, the location and class of each detected patch is saved for all the images in the dataset along with the confidence.

## 3 Experimental Setup, Datasets, and Evaluation Criteria

We use two image datasets, one fluorescence microscopy and one immunohistochemistry to quantitatively evaluate the performance of SimSearch. Two other datasets, one transmission electron microscopy (TEM) and one blood smear brightfield microscopy dataset are used for ablation studies to evaluate the influence of different design choices and parameter settings in the framework.

### 3.1 Cell Classification for GFP Translocation Analysis

The first evaluation dataset consists of fluorescence microscopy images from the open dataset originally provided by Ilya Ravkin and made publicly available via the Broad Bioimage Benchmark Collection (15). The images are from a drug screening experiment, where human U2OS cells are grown in a 96-well plate with varying dose of two drugs. As the drug dose increases, a protein tagged with the green fluorescent protein GFP is translocated from the cytoplasm to the nucleus, and thus the amount of GFP expressed in the nuclei increases and GFP expressed in the cytoplasms decreases. The goal of the analysis is to quantify this translocation of GFP, or more specifically, to measure the fraction of cells in an image that have nuclear or cytoplasmic GFP expression. Not all cells express GFP, meaning that we also need a third class, referred to as the “no GFP” class. This gives three classes of interest for the experiment, i.e, “GFP in Nucleus”, “GFP in Cytoplasm”, and “No GFP”.

The experimental setup using SimSearch consists of a workflow where the user is asked to define the three classes of cells as described above, and the output is the number of patches per image representing the three classes. Five *prototype* patches of size 48 ×48 pixels from each of the “GFP in Nucleus”, “GFP in Cytoplasm”, and “No GFP” were selected from six images and the threshold was adjusted individually until five *negative* patches were marked for each class. The confidence threshold was set to 0.65 for each class after manual inspection of the results.

Here we compare the results achieved with SimSearch to those achieved with CellProfiler (3) and CellProfilerAnalyst(4). We tuned a CellProfiler pipeline on the translocation dataset, and extracted a large number of intensity and morphology features. We thereafter trained a RandomForest classifier in CellProfilerAnalyst to detect the three cell classes described above. In order to make the comparison between SimSearch and CellProfiler Analyst as fair as possible, we used the same set of training cells as were used in the first step of the SimSearch workflow.

To evaluate the SimSearch performance, we compared the classification results to those obtained using CellProfiler/CellProfilerAnalyst in relation to the known drug dose per well. The ratio of cells from each class with respect to the total number of cells detected in each image in CellProfiler and the ratio of patches of each class with respect to all the patches detected in SimSearch was calculated and plotted against the drug dose.

### 3.2 Segmentation of Brightfield Images of Tissue

The second evaluation dataset consists of publicly available tissue samples obtained from the Human Protein Atlas (16), where protein expression in different human tissue types is derived from antibody-based immunohistochemistry. Antibodies are labeled with DAB (3,3’-Diaminobenzidine) and the resulting brown stain indicates where an antibody has bound to its corresponding antigen. The sections are furthermore counterstained with hematoxylin to enable visualization of other tissue components, such as cell nuclei. Ethical approval was not required for usage of this human data as confirmed by the license attached with the open access data.

In our experimental setup, we wanted to investigate SimSearch’s ability to identify different sub-cellular protein localizations, and therefor chose two proteins with known localizations: ERBB2, which is membranous and BRCA1 which is located mostly in nuclei. To limit the influence of variation in general tissue morphology, we included (when available) paired ERBB2 and BRCA1 samples from the same patient. Note that all tissue samples also consist of various amounts of tissue without DAB stain. In total 33 images were extracted and used in the study.

Five *prototype* patches of 48 × 48 pixels each representing nuclear stain, membranous stain, and tissue without DAB stain were selected from six training images. The similarity threshold was decreased individually until a representative number of patches were displayed in the images, and approximately ten *negative* patches were marked for each class. Once trained, the classifier was applied to the full dataset. The confidence threshold was set to be 0.6 for each class. This was motivated by the need to get a more exact outline of the different regions in the tissue, and a more stable quantification of the different tissue fractions.

For this dataset, the output from SimSearch was modified to provide a semantic segmentation of the tissue sample. This was done by resizing the patchwise prediction outputs to the original image size, and assigning the class with maximum probability above the confidence threshold at each pixel as final prediction.

### 3.3 Ablation Studies

Ablation studies were performed to assess the variability in performance of the framework under different training conditions. Datasets of transmission electron microscopy images and blood smear bright-field microscopy images were used for this purpose.

#### Cilia Detection

The first ablation dataset consists of negative stain transmission electron microscopy (TEM) images of cell sections for cilia morphology analysis. Mouth swabs of respiratory epithelial cells of which some have protruding cilia were redissolved, fixated, and embedded in plastic before sectioned into slices of (≈50-70 nm) in thickness, using a microtome. The slices of the specimen were then floated onto carbon coated copper mesh grids and post-stained to increase contrast. The images were acquired with a low-voltage (25keV) MiniTEM (Vironova AB, Stockholm, Sweden) as 16-bit TIFF images of size 2048×2048 pixels and a pixel size of 1nm. The cilia positions were annotated using bounding boxes and the centers were treated as the ground truth. The dataset consisted of 39 images with a total of 393 cilia, of which 31 images with 296 cilia were used for training and eight images with 97 cilia were kept separately for testing. From the training set, six random images were displayed and used for user interaction, and the rest were used for noisy data extraction. The performance was evaluated on the test set. The patch size was chosen to be 72 × 72 pixels to include also slightly bigger (stretched during sectioning) cilia as well.

#### Trophozoites and RBC Detection

The other dataset used for ablation studies consists of image set BBBC041v1 made publicly available via Broad Bioimage Benchmark Collection (15). The dataset consists of 1384 brightfield microscopy images of blood smears stained with Giemsa reagent divided into three different sets, containing two classes of uninfected cells (RBCs and leukocytes) and four classes of infected cells (gametocytes, rings, trophozoites, and schizonts). For the purpose of this study, a subset of the images containing only RBCs and trophozoites was used as the other classes were very rare. This meant that 258 images containing 607 trophozoites and 19784 RBCs were used in the experiments.

Of these, 206 images with 502 trophozoites and 15262 RBCs were used for training and 52 images with 105 trophozoites and 4522 RBCs were kept separate for testing. Ten random images from the training set were chosen to be displayed for user interaction, and the rest of the images in the training set were used for noisy data extraction. The performance was evaluated on the test set. To utilize the iterative patch selection better in this evaluation study, the random image set chosen for display were required to have at least 20 trophozoites present in the images. The patch size was chosen to be 64 × 64 pixels.

In both ablation studies, a detected patch is counted as a true positive (TP) if the distance between the center of the ground truth patch and detected patch is less than three times the stride of the sliding window patch extraction. The rest of the detected patches are counted as false positives (FPs) and the undetected ground truth patches are counted as false negatives (FNs).

The performance of the framework was measured on the area under the precision-recall curve (AUC) by varying the number of *prototypes* and *negative* patches. The number of *prototype* patches *P* was varied as *P* = {5, 10} and the number of *negative* patches *N* was varied as *N* = {0, 5}. This was achieved by selecting P ground truth patches randomly and then adjusting the similarity threshold until only *N negative* patches remained in the displayed images. The effect of different training strategies, especially the self-supervised training on noisy patches and negative supervised contrastive training which were introduced in a novel way in the framework was also compared. *Pos* training refers to training only with the *clean positive* patches obtained during the iterative patch selection, *Pos* + *Neg* training refers to the training with both *clean positive* and *clean negative* patches, and *Selflearn* refers to the self-supervised training of the *noisy* patches as described in Section 2.3.3.

For statistical significance, the experiments in each combination of *P* and *N* were repeated 20 times by selecting new images and new prototypes for iterative patch selection randomly. Since the user-approved data is minimal (5-15 patches per class), a validation set of patches to assess model performance cannot be meaningfully extracted without affecting the performance. Currently, the training is stopped after a fixed number of iterations (empirically chosen) which results in some variance in the results. Hence, to test the repeatability of a particular training strategy, each training strategy was repeated five times (with the same training data and hyper-parameter settings) for each iteration and the standard deviation in AUC is recorded.

In the first experiment, the effect of the number of *P* and *N* on the AUC was computed. In the second experiment, the effect of training strategies on the AUC was compared. Lastly, the repeatability of the training strategies was shown by computing the mean standard deviation in AUC for a particular training strategy.

All the strategies were compared against the fully supervised baseline using *Pos* training on the whole training set to qualitatively assess the performance of SimSearch with the fully supervised approach that requires significantly larger sets of labelled data. The background patches were extracted from images by removing the labelled class patches. For supervised baseline training in cilia experiments, 311 patches were extracted, and for trophozoites and RBCs experiments, 913 patches were extracted. The supervised experiments were also repeated 20 times and mean and standard deviation of the AUC was recorded.

### 3.4 Implementation Details

#### Iterative Patch Selection

In all experiments, patch selection was done based on the features extracted from the pre-trained ResNet-18 (17) network trained on ImageNet (11). Since the deeper layers are task specific, only the first 3 blocks of ResNet were used. A convolutional filter of size (*H*/16 × *W*/16), where *H* and *W* are the height and width of the patch respectively, initialized with a gaussian was added to get a center focused single feature vector output for every patch. The inference was done on NVIDIA Titan X GPU with 12 GB and took approx. 0.5 sec/image for initial patch feature extraction (which happens when the first patch is selected) and SE is almost instantaneous (milliseconds) afterwards. The generation of noisy training patches also takes approx. 0.5 sec/image.

#### Noisy Training

For self-supervised training, a projection head consisting of 2 fully-connected layers was added. The model architecture is shown in Figure 3. The initial convolution layer and the first 2 blocks of ResNet were frozen and the network was then trained for 100 epochs with the Adam optimizer (18) and a learning rate of 10^-3^ with a step decay of 0.05 after each 50 epochs. The model was trained with a batch size of 128 and augmentations were performed with random horizontal and vertical flips, rotation, solarize, coarse-dropout, and brightness and contrast changes. *τ_self_* was chosen to be 0.5.

#### Supervised Refinement

For supervised contrastive training, the model is then trained with a batch size of 32 for 50 epochs with Adam optimizer and a learning rate of 10^-3^ with a step decay of 0.1 after each 25 epochs. *τ_sup_* was chosen to be 0.07. The same augmentation strategy as in the self-supervised training step except solarize and coarse-dropout was used.

## 4 Results and Discussion

### 4.1 Cell Classification for GFP Translocation Analysis

As shown in Figure 4, the output of SimSearch corresponds well with the drug dose in the wells. The fraction of ROIs with GFP expression in the nuclei increases sharply as the drug dose reaches 15.6 nM. Similarly, the fraction of ROIs with GFP expression in the cytoplasms decreases at the same drug dose. The fraction of ROIs of cells with no GFP (which is independent of drug dose) remains similar. At each dose, the SimSearch results follow the CellProfiler results very well. However the magnitude differs slightly. This is likely due to the fixed bounding box size in SimSearch, i.e., if cells of the same class are close together, they will be detected and counted as one. As a result, the ratio of cells with GFP in cytoplasm is higher and No GFP is lower for smaller drug doses in SimSearch.

**Figure 4:**
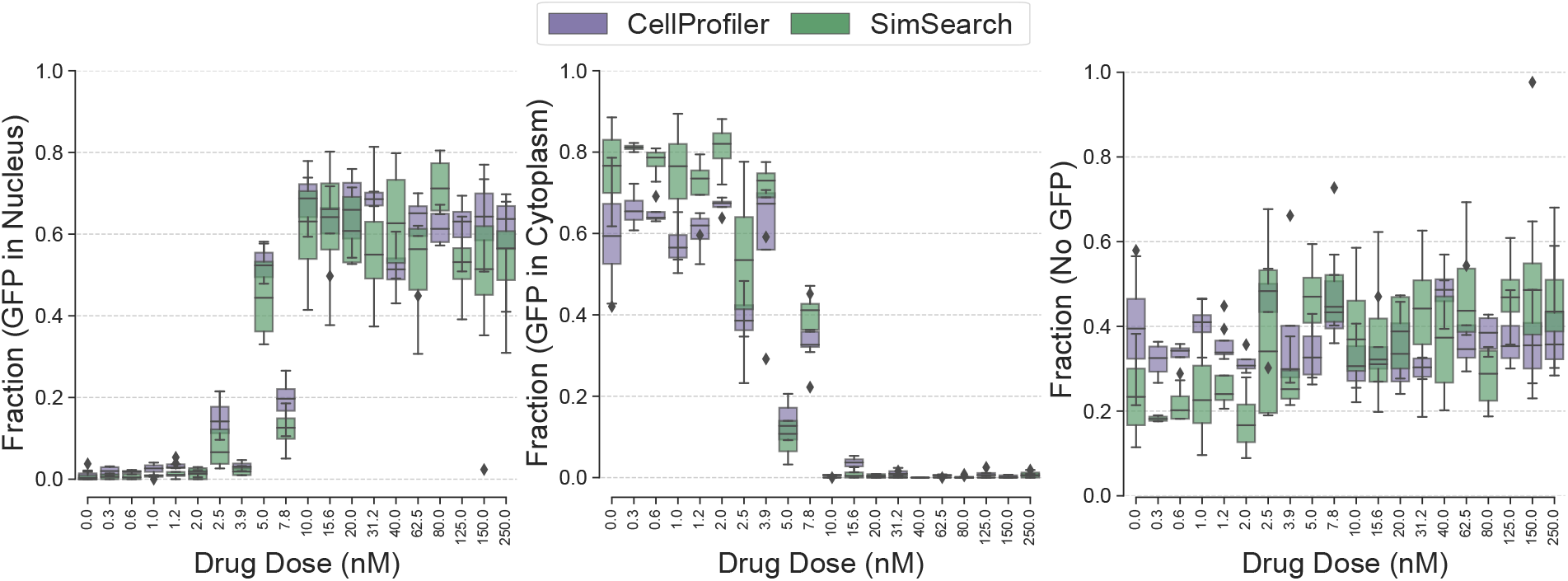
Comparison between CellProfiler pipeline and SimSearch results. The fraction of each cell class of the total number of cells detected is shown with respect to the drug dose.

As can be seen in Figure 5, the detected ROI centers are usually not located perfectly at the center of the cells. The objects are not symmetrical, and the *clean* ROIs detected in the iterative patch selection process are compared to the *prototype* patches and their rotations. The similarity local maximum then falls to the center of the patch and not the center of the object. These patches are then used in training the model, and hence the local maxima found in the final detections are usually at a small distance from the corresponding cell centers.

**Figure 5:**
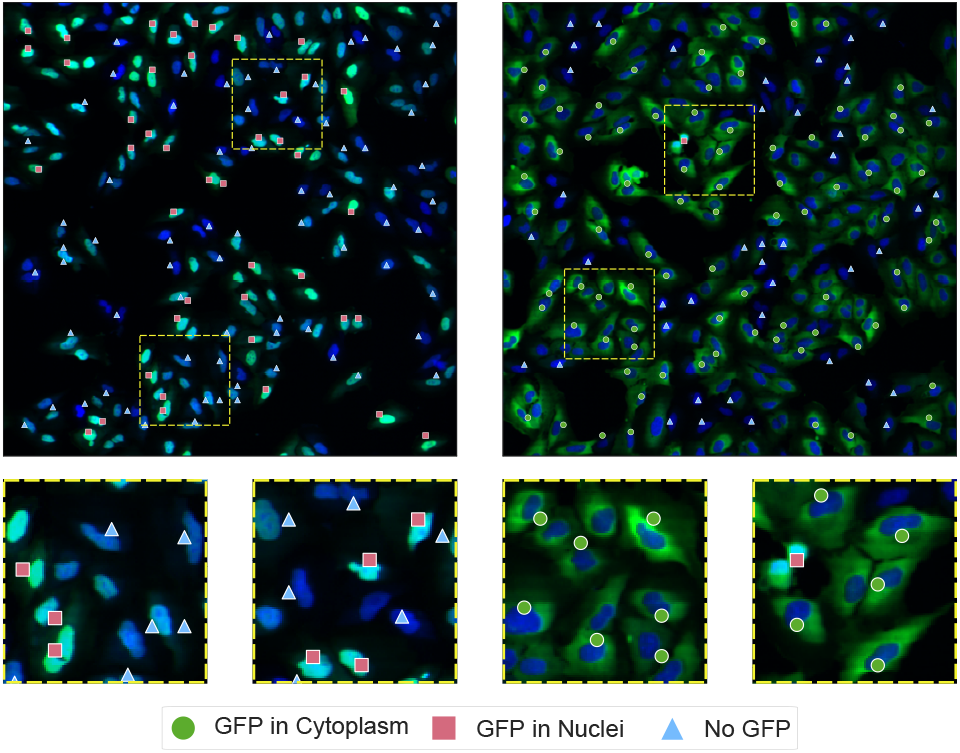
Display of the SimSearch results on the GFP translocation images. The centers of the detected ROIs of the different classes are marked.

### 4.2 Segmentation of Brightfield Images of Tissue

The area fraction of tissue classified by SimSearch as showing membraneous and nuclear DAB-staining patterns, as well as no DAB stain is presented in Figure 6. The area fraction classified as having membranous DAB is generally higher in samples stained for ERBB2, while the area fraction with nuclear stain is higher in samples stained for BRCA1, as expected. Examples of the full semantic segmentation results are shown in Figure 7. SimSearch is fairly consistent in identifying membraneous and nuclear stain localization, as illustrated by the examples A,B, E, and F in the top half of Figure 7. The bottom half of the figure shows some difficult examples. C shows an example where the ERBB2 staining (which should be membraneous) is so strong it appears to also stain nuclei. Similarly, G shows a sample stained for BRCA1, which should be nuclear, but SimSearch finds both nuclear and membraneous patterns. Careful inspection reveals that the DAB staining has spread to the surrounding tissue, resulting in a false membranous pattern.

**Figure 6:**
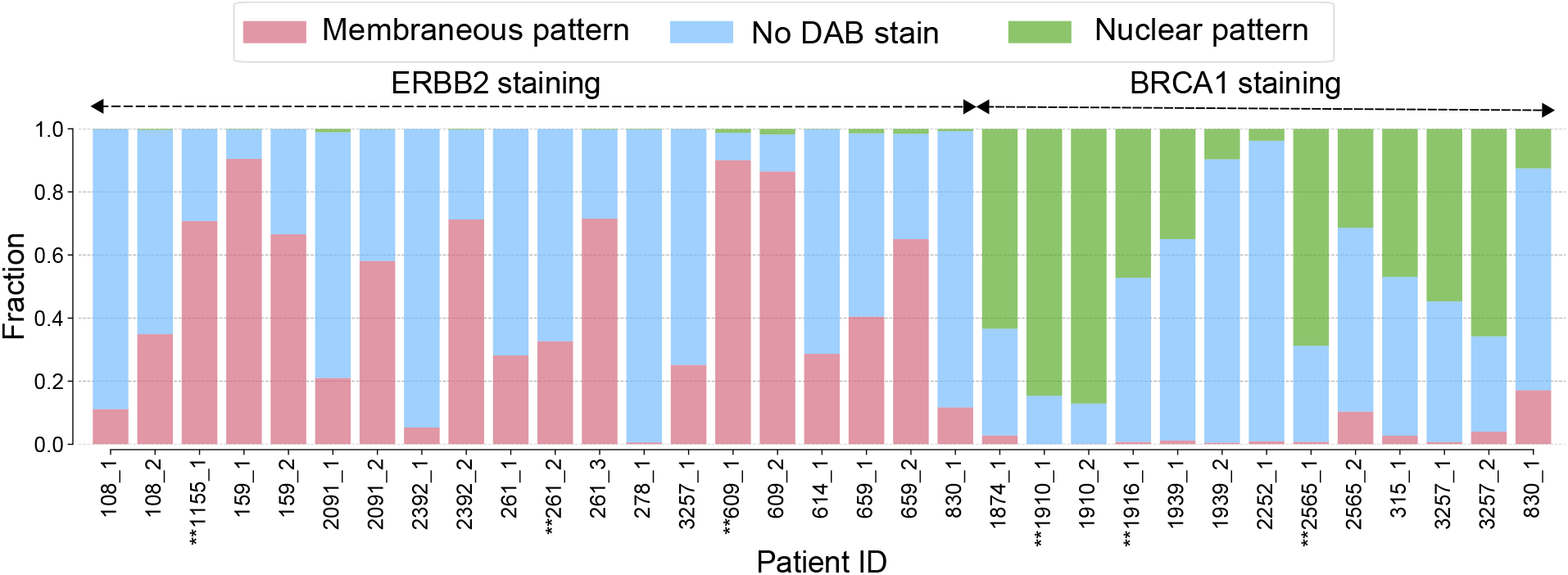
Fraction of tissue classified as containing membraneous, nuclear, or no DAB-staining patterns, correlating with targeted proteins ERBB2 and BRCA1. Images marked with an asterisks were used during training.

**Figure 7:**
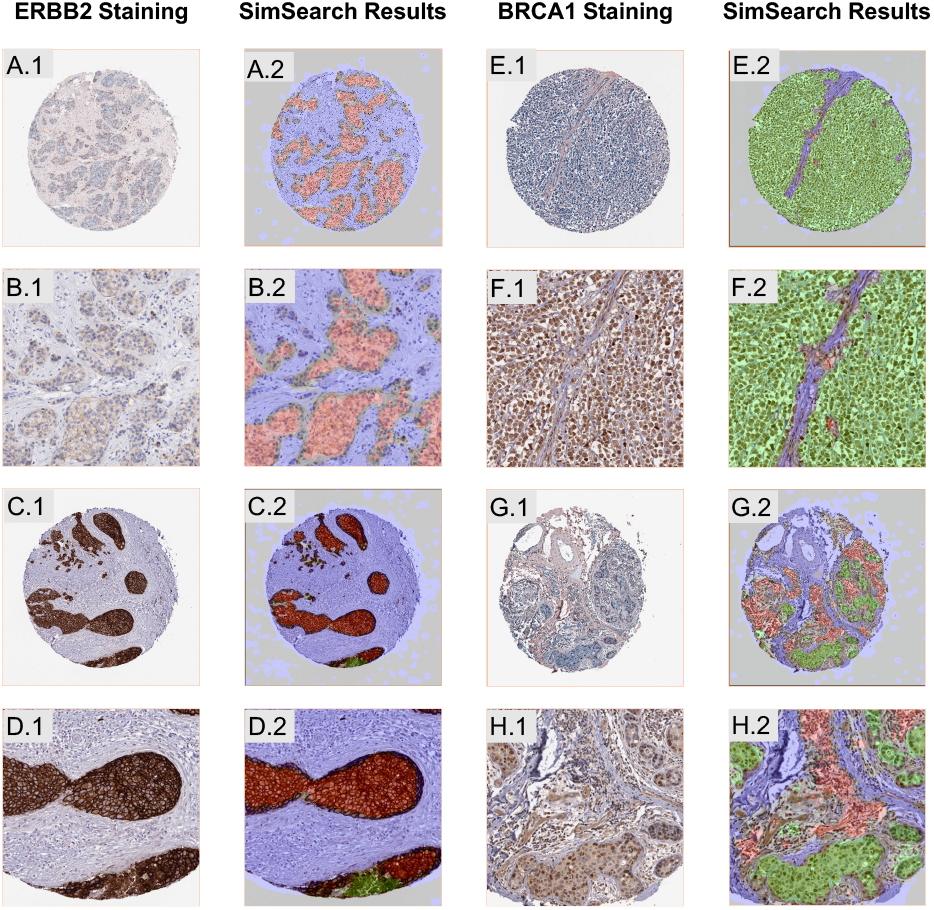
Examples of DAB staining patterns and the resulting patch classification and semantic segmentation by SimSearch. **A-D** ERBB2 stained tissue cores. **E-H** BRCA1 stained tissue cores. Regions classified as having membraneous patterns are overlaid with red, while nuclear DAB patterns are overlaid with green, and regions without DAB stain are overlaid in blue. B, D, F, and H show a zoomed-in region of A, C, E, and G respectively. Note that C.2 and G.2 show mixed classification results.

### 4.3 Ablation Studies

Figures 8 and 9 show the results of the ablation studies on the Cilia and Blood Smear datasets respectively. The violin plots show the probability density of the data at different values. The “body”, hence, shows where the values are concentrated for different experiments and the long tail with the neck shows the outliers. The normal box plot is shown inside the violin.

**Figure 8:**
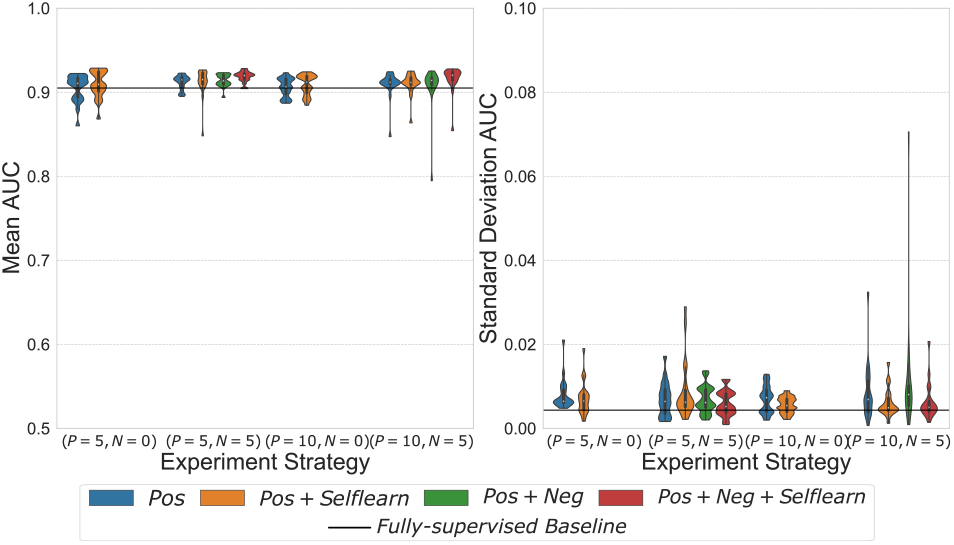
Area under precision-recall curve (AUC) comparison of different training strategies and number of training examples on the Cilia dataset. On the left, we show the mean AUC (of 5 iterations) of each training strategy for 20 experiments, and on the right, the standard deviations of the iterations for each experiment. The mean AUC and standard deviation of 20 repetitions of fully supervised baseline is shown as horizontal lines in the plots.

**Figure 9:**
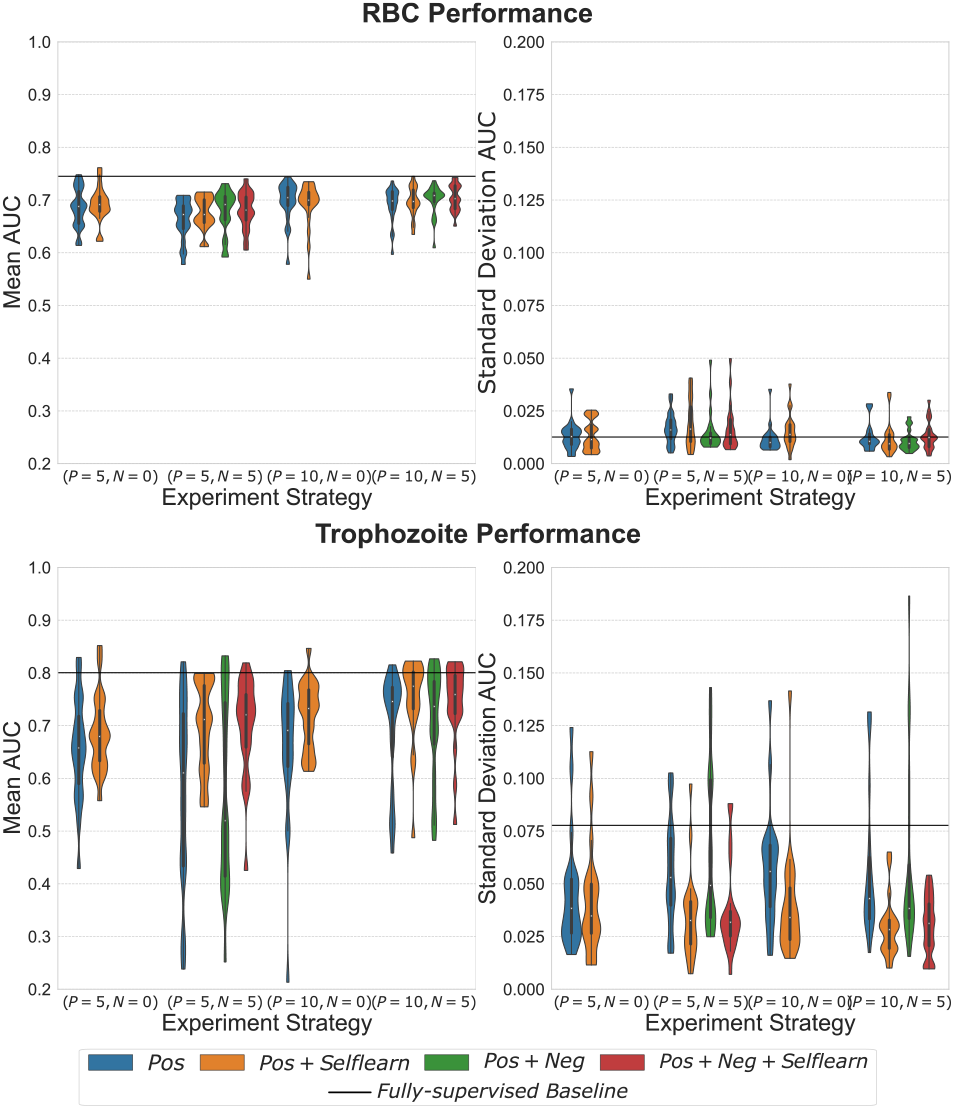
Area under precision-recall curve (AUC) comparison of different training strategies and number of training examples on the blood smear dataset. On left (top and bottom), mean AUC (of 5 iterations) of the each training strategy for 20 experiments (for RBCs and Trophozoites respectively). On right, the standard deviations of the iterations for each experiment (for RBCs and Trophozoites respectively). Mean AUC and standard deviation of 20 repetitions of fully supervised baseline is shown as horizontal lines in the plots.

The left plots of Figures 8 and 9 show mean AUC (of 5 repetitions) for 20 different experiments of each combination of number of *prototypes*(*P*) and *negatives*(*N*). This plot shows how the AUC varies with the different numbers of *prototypes* and *negatives* and training strategies. It also shows how different training strategies adapt with different selections of *prototypes* and *negatives* in a category.

The right plots of Figures 8 and 9 show the standard deviation of AUC (of 5 repetitions) for the different *P* and *N* combinations and training strategies. This plot shows how stable a particular training strategy is, given a *clean* and *noisy* dataset and helps in accessing the repeatability of an experiment. Higher repeatability corresponds to lower variance for a training scheme.

As can be seen from Figures 8 and 9 (left), there is not a significant difference in the median AUC for different combinations of *P* and *N*, which shows that even with a smaller number of samples a reasonable performance can be achieved. However, the necks become shorter with an increase in *P* (except for training with only positives). This indicates that performance is less dependent on selecting a good *prototype* if more examples of both positives and negatives are included. A similar effect can be seen with the introduction of self-learning within the same *P, N* category (smaller body and shorter necks) which indicates that self-learning helps make the AUC more robust with respect to the choices of random patches in iterative patch selection process.

As can be seen in Figures 8 and 9 (right), self-learning makes the framework more robust in general (as indicated by smaller standard deviation between repetitions of the same experiment), thus increasing repeatability and making the results more reliable.

In comparison with the fully supervised approach, as can be seen from Figure 8, for cilia most combinations of *P* and *N* outperform the baseline, however, the standard deviation seems to be higher than the baseline. Similarly, for the RBC and trophozoite experiments, even with relatively low numbers of *P* and *N* the performance can reach the mean of the supervised baseline and even surpass it in some cases. It is also worth noting that the standard deviation is usually lower than that of the supervised baseline which shows the robustness of the method. This clearly shows the advantages of using SimSearch over manually labelling a large dataset.

In the case of abundant and “easy” classes like Cilia and RBCs (as the objects in these classes have definitive shape with small variations in size and color), training with *negative* patches (*Pos* + *Neg*) helps reduce the inter-measurement variability (lower standard deviation between repeated measurements). It also makes the model more independent of the choice of *prototypes* and reduces variance in mean AUC between different experiments. *Pos* + *Neg* training helps the model learn more relevant features for a class which help to distinguish it from other classes.

However, for a rare and “difficult” class like Trophozoites (objects with large variation in shape, size and color), training with just (*Pos* + *Neg*) does not seem to improve performance. This can be attributed to fewer *clean positive* patches being available for the model to learn the relevant features for the class. However, self-learning with noisy patches in this case increases the performance significantly. This shows the effectiveness of self-learning in learning the general features for all the classes. This is shown in reduction of both inter-experiment (Figure 9(bottom left)) and inter-measurement (Figure 9 (bottom right)) standard deviation. Although, *Pos* + *Selflearn* has higher median than *Pos* + *Neg + Selflearn* for trophozoites, overall *Pos + Neg + Selflearn* seems to be the better choice for both the “easy” and “hard” classes in a dataset.

The numerical results of comparison between different experiment and training strategy is available in Supplementary Table I, II, and III for Cilia, RBCs and Trophozoites respectively.

## 5 Conclusions

We have presented a human-in-the-loop framework for fast and flexible ROI detection in biological image datasets. The proposed framework uses a pretrained model to extract features removing the need for feature engineering and domain-expertise from the user. The framework is fast when using a GPU which makes it suitable for real-time applications. The framework employs self-supervised learning and negative training to make the most efficient use of user input during the training process. The GUI also provides the user with a confidence threshold to control the output of the experiments in accordance with what is most relevant/important for the research question/application at hand. We demonstrated the framework on different research scenarios and four biological datasets, and presented good performance and robustness under different prerequisites and requirements. The framework successfully reciprocated the results from CellProfiler in drug response analysis without manual and time-consuming feature selection and extraction. The framework also performed well detecting and segmenting areas exposed to different immunohistochemical stains. Using ablation experiments, we showed that the training strategy and methods implemented in the framework are robust against different variations in user inputs. We hence conclude that we have shown that the framework has great potential to increase research throughput in a broad variety of biological microscopy experiments.

## References

[1] C. T. Rueden, J. Schindelin, M. C. Hiner, B. E. DeZonia, A. E. Walter, E. T. Arena, and K. W. Eliceiri, “Imagej2: Imagej for the next generation of scientific image data,” BMC bioinformatics, vol. 18, no. 1, pp. 1–26, 2017.

[2] J. Schindelin, I. Arganda-Carreras, E. Frise, V. Kaynig, M. Longair, T. Pietzsch, S. Preibisch, C. Rueden, S. Saalfeld, B. Schmid et al., “Fiji: an open-source platform for biological-image analysis,” Nature methods, vol. 9, no. 7, pp. 676–682, 2012.

[3] C. McQuin, A. Goodman, V. Chernyshev, L. Kamentsky, B. A. Cimini, K. W. Karhohs, M. Doan, L. Ding, S. M. Rafelski, D. Thirstrup et al., “Cellprofiler 3.0: Next-generation image processing for biology,” PLoS biology, vol. 16, no. 7, p. e2005970, 2018.

[4] D. Dao, A. N. Fraser, J. Hung, V. Ljosa, S. Singh, and A. E. Carpenter, “Cellprofiler analyst: interactive data exploration, analysis and classification of large biological image sets,” Bioinformatics, vol. 32, no. 20, pp. 3210–3212, 2016.

[5] S. Berg, D. Kutra, T. Kroeger, C. N. Straehle, B. X. Kausler, C. Haubold, M. Schiegg, J. Ales, T. Beier, M. Rudy et al., “Ilastik: interactive machine learning for (bio) image analysis,” Nature Methods, pp. 1–7, 2019.

[6] A. Gupta, P. J. Harrison, H. Wieslander, N. Pielawski, K. Kartasalo, G. Partel, L. Solorzano, A. Suveer, A. H. Klemm, O. Spjuth et al., “Deep learning in image cytometry: a review,” Cytometry Part A, vol. 95, no. 4, pp. 366–380, 2019.

[7] E. Moen, D. Bannon, T. Kudo, W. Graf, M. Covert, and D. Van Valen, “Deep learning for cellular image analysis,” Nature methods, pp. 1–14, 2019.

[8] Y. LeCun, Y. Bengio, and G. Hinton, “Deep learning,” nature, vol. 521, no. 7553, pp. 436–444, 2015.

[9] L. Jing and Y. Tian, “Self-supervised visual feature learning with deep neural networks: A survey,” IEEE Transactions on Pattern Analysis and Machine Intelligence, pp. 1–1, 2020.

[10] X. Liu, F. Zhang, Z. Hou, L. Mian, Z. Wang, J. Zhang, and J. Tang, “Self-supervised learning: Generative or contrastive,” IEEE Transactions on Knowledge and Data Engineering, 2021.

[11] J. Deng, W. Dong, R. Socher, L.-J. Li, K. Li, and L. Fei-Fei, “Imagenet: A large-scale hierarchical image database,” in 2009 IEEE conference on computer vision and pattern recognition. Ieee, 2009, pp. 248–255.

[12] T. Chen, S. Kornblith, M. Norouzi, and G. Hinton, “A simple framework for contrastive learning of visual representations,” in International conference on machine learning. PMLR, 2020, pp. 1597–1607.

[13] J. Han, P. Luo, and X. Wang, “Deep self-learning from noisy labels,” in Proceedings of the IEEE International Conference on Computer Vision, 2019, pp. 5138–5147.

[14] P. Khosla, P. Teterwak, C. Wang, A. Sarna, Y. Tian, P. Isola, A. Maschinot, C. Liu, and D. Krishnan, “Supervised contrastive learning,” arXiv preprint arXiv:2004.11362, 2020.

[15] V. Ljosa, K. L. Sokolnicki, and A. E. Carpenter, “Annotated high-throughput microscopy image sets for validation.” Nature methods, vol. 9, no. 7, pp. 637–637, 2012.

[16] M. Uhlén, L. Fagerberg, B. M. Hallström, C. Lindskog, P. Oksvold, A. Mardinoglu, Å. Sivertsson, C. Kampf, E. Sjöstedt, A. Asplund et al., “Tissue-based map of the human proteome,” Science, vol. 347, no. 6220, 2015.

[17] K. He, X. Zhang, S. Ren, and J. Sun, “Deep residual learning for image recognition,” in Proceedings of the IEEE conference on computer vision and pattern recognition, 2016, pp. 770–778.

[18] D. P. Kingma and J. Ba, “Adam: A method for stochastic optimization,” arXiv preprint arXiv:1412.6980, 2014.

